# SingleChannelNet: A Model for Automatic Sleep Stage Classification with Raw Single-Channel EEG

**DOI:** 10.1101/2020.09.21.306597

**Authors:** Dongdong Zhou, Guoqiang Hu, Jiacheng Zhang, Jian Wang, Rui Yan, Fan Li, Qi Xu, Lauri Kettunen, Zheng Chang, Fengyu Cong

## Abstract

In diagnosing sleep disorders, sleep stage classification is a very essential yet time-consuming process. Most of the existing state-of-the-art approaches rely on hand-crafted features and multi-modality polysomnography (PSG) data, where prior knowledge is compulsory and high computation cost can be expected. Besides, few studies are able to obtain high accuracy sleep staging using raw single-channel electroencephalogram (EEG). To overcome these shortcomings, this paper proposes an end-to-end framework with a deep neural network, namely SingleChannelNet, for automatic sleep stage classification based on raw single-channel EEG. The proposed model utilizes a 90s epoch as the textual input and employs two multi-convolution blocks and several max-average pooling layers to learn different scales of feature representations. To demonstrate the efficiency of the proposed model, we evaluate our model using different raw single-channel EEGs (C4/A1 and Fpz-Cz) on two different datasets (CCSHS and Sleep-EDF datasets). Experimental results show that the proposed architecture can achieve better overall accuracy and Cohen’s kappa (CCSHS: 90.2%-86.5%, Sleep-EDF: 86.1%-80.5%) compared with state-of-the-art approaches. Additionally, the proposed model can learn features automatically for sleep stage classification using different single-channel EEGs with distinct sampling rates from different datasets without using any hand-engineered features.

## I. Introduction

**H**UMANS spend about one-third time of life on sleeping, and high-quality sleep plays a vitally important role in the restoration of body and mind [1]. Whereas roughly 33% of the population in the world suffers from insomnia disorder [2]. Correctly identifying sleep stage using whole-night PSG data is essential to diagnose and treat sleep-related disorders [3]–[6]. The PSG recordings comprise of the EEG, electrocardiogram (ECG), electrooculogram (EOG), electromyogram (EMG) and other respiration signals [7].

According to the guidelines of the Rechtschaffen and Kales (R&K) [8] or American Academy of Sleep Medicine (AASM) [9], the PSG data should be first segmented into 30s epochs typically, then these sequential epochs are defined as different stages. Some sleep-related disorders have particular sleep structure, it is therefore beneficial to diagnose them with accurate sleep stage classification. Traditionally, the sleep stage classification task is conducted by experts manually following the R&K and AASM rule which is often time-consuming, labor-intensive and prone to subjective mistakes [6]. Hence, there is an urgent need for automatic sleep stage classification approach to assist the clinician’s work and achieve reliable results.

Some methods based on machine learning have been proposed to identify the sleep stage. These approaches generally extract either time-domain features [3], [10], [11] or frequency-domain features [12]–[16] from the PSG signals and these pre-extracted features are then fed into the conventional classifier, such as support vector machine (SVM) [4], [14], [17], [18], *k*-nearest neighbors (KNN) [16], [19], [20], random forest [21]–[24] and so on. The performance tremendously relies on the categories and the number of features, which are extracted based on the characteristics of experimental datasets. Therefore, these approaches may not be robust enough to be generalized to different datasets because of the distinct properties between datasets.

In recent years, the deep networks show great capacity for automatic features learning from data, and it can avoid the reliance on hand-engineered features. Meanwhile, a series of deep learning methods are applied to sleep stage classification. Here, we categorize these approaches into multi-channel [6], [25]–[29] versus single-channel schemes [30]–[35] based on the number of input channels. Following the multi-channel scheme, Phan *et al*. [6] first transformed the raw signals into the time-frequency image through the short-time Fourier transform as the input of the proposed convolutional neural network (CNN). The overall accuracy achieved was equal to 82.3%, in which there is room for improvement. Besides, the time-frequency image relies much on many preprocessing steps, it would be time-consuming and in need of prior knowledge of signal processing. Aiming at this, Chambon *et al*. [27] proposed a novel network architecture of low computational cost adopting multivariate and multimodal time series from EEG, EMG and EOG, but the classification performance is not good enough with the accuracy of 80% compared to state-of-the-art methodologies. One important reason is that the convolutional layers with fixed filter size were stacked sequentially, which can not learn multiscale features simultaneously. A promising approach was proposed by Zhang *et al*. [29], who employed the CNN and recurrent neural network (RNN) to capture temporal and spatial information simultaneously from the PSG data. The architecture attained an accuracy of 87%. Although the combination of CNNs and RNNs can enhance the model performance to some extent, the high computational cost of RNNs should be taken into consideration. To the best of our knowledge, the training speed of CNNs would be dozens of times faster than that of RNNs under the same GPU acceleration when implementing long time-series input. To sum up, despite the fact that multi-channel PSG data can provide additional referenced information compared to single-channel EEG, there is also some irrelevant information being introduced. Furthermore, multi-channel recordings can limit the practical application on account of more complex operation and equipment costs.

Compared to the multi-channel scheme, the single-channel scheme can reduce the related cost and be much easier for data acquisition. Under the single-channel scheme, Supratak *et al*. [30] introduced a deep learning model called DeepSleepNet. DeepSleepNet utilizes the capacity of deep learning to extract time-invariant features automatically, the proposed model can be adapted to different datasets. However, the accuracy obtained from DeepSleepNet was 82%, which can not outperform the state-of-the-art approaches. A promising CNN model was proposed by Sors *et al*. [31], who used raw single-channel EEG to classify the sleep stage without any preprocessing. The architecture attained an accuracy of 87%, whereas the model complexity is a bit high with 12 convolutional layers. Furthermore, the filer size was chosen among 7, 5, 3, the performance of larger size filters should be compared considering the long length of input (1.5 × 10^4^).

To tackle these problems, this paper proposes the SingleChannelNet (SCNet), a model for automatic sleep stage classification based on raw single-channel EEG, which can learn different scales features simultaneously. We aim to automate the sleep stage classification completely by utilizing the capabilities of the proposed model. The main contributions of this work are as follows:

i. We propose a new deep learning model with low model complexity for sleep stage classification using 90s raw single-channel EEG.
ii. We implement two multi-convolution (MC) blocks with different filter sizes in our model. In addition, the max-average (M-Apooling) layer is applied to take place of the conventional max-pooling layer. Two strategies are used for capturing more feature representations from different scales to enhance the capacity of the feature extraction.
iii. The results demonstrate that our model can obtain promising performance on different raw single-channel EEGs (C4/A1, Fpz-Cz) from CCSHS and Sleep-EDF datasets, without modifying the architecture and hyper-parameters of model and training algorithm. Moreover, all features are learned by the proposed model automatically.

The rest of this paper is organized as follows. We represent the experimental datasets in Sec. II. Sec. III describes the structure of the SCNet model and the training algorithm. In Sec. IV, the experimental results are represented. The final discussion and conclusion are included in Sec. V.

## II. Experimental Datasets

Two public datasets are employed to evaluate the performance of the proposed framework in this work, namely Cleveland Children’s Sleep and Health Study (CCSHS) [36], [37] and Sleep-EDF Database Expanded (Sleep-EDF, 2018 version) [38]. It should be noted that all hypnograms of experimental datasets are manually scored according to the R&K manual rather than the AASM rule.

### A. Cleveland Children’s Sleep and Health Study (CCSHS)

The CCSHS dataset comprises of overnight PSG recordings from 515 subjects aged 8-11 years, which is one of the largest population-based pediatric cohorts studied with objective sleep studies. Each 30s epoch is manually divided by experts into several stages: Wake (W), Rapid Eye Movement (REM), Non-REM1 (N1), Non-REM2 (N2), and Non-REM3 (N3). In this work, single-channel EEG C4/A1 sampled at 128 Hz is selected.

### B. Sleep-EDF Database Expanded (Sleep-EDF)

The Sleep-EDF dataset consists of two subsets: sleep-cassette (SC) contains 78 healthy Caucasians aged from 25 to 101 years and sleep-telemetry (ST) comprises 22 Caucasians receiving temazepam treatment. Each participant was recorded two subsequent night PSG data except the subject 13, subject 36 and subject 52, from the SC subset who had only a one-night record. Each epoch of recordings is manually labelled by clinicians according to the R&K rule into W, N1, N2, N3, N4, REM, MOVEMENT and UNKNOWN stages respectively.

In addition, MOVEMENT and UNKNOWN are excluded, as they do not belong to the six stages. The PSG data include two-channel EEGs (Fpz-Cz and Pz-Oz), single-channel EOG, single-channel EMG and the event marker (sampled at 1 Hz). The sampling rate *f*_*s*_ of EEG, EOG, and EMG is 100 Hz. Single-channel EEG Fpz-Cz is adopted in our experiment. For the Sleep-EDF dataset, stages N3 and N4 are merged into stage N3 which is consistent with the AASM manual. Additionally, resampling operation is not applied to C4/A1 and Fpz-Cz EEGs. We also found that most previous studies use the Sleep-EDF dataset of the first 20 subjects (Sleep-EDF-v1). For a fairer comparison, we also experiment with the Sleep-EDF-v1 dataset.

### C. Contextual input

In previous works, most schemes use a single 30s epoch as the classifier input [7], [31], [35] and then produce a single output label. Although being straightforward, this classification method ignores the existing correlation and dependency between surrounding epochs. It is considered that the sleep stage classification depends not only on the local epoch, but also on the prior and following temporal features [6], [9]. For this reason, an extension of single 30s epoch input is conducted by combining it with its neighboring epochs to make a contextual input. Furthermore, we employ 90s epoch (**Z**_*m*_) as contextual input of the proposed model, and it contains three sequential epochs: prior 30s epoch (**X**_*m−*1_), 30s epoch (**X**_*m*_) and subsequent 30s epoch (**X**_*m*+1_). The ground truth label of **Z**_*m*_ is *y*_*m*_ which also denotes **X**_*m*_’s label. As in

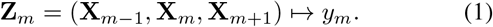

Details are illustrated in Fig. 1. As shown in Table I, we summarize the number of 90s epochs for each sleep stage from CCSHS, Sleep-EDF and Sleep-EDF-v1 datasets in our experiments. The distribution of the number of five stages is imbalanced. For all datasets, W and N2 stages account for more than 60% of all 90s epochs. By contrast, the proportion of stages N1 and N3 is the smallest.

**Fig. 1.**
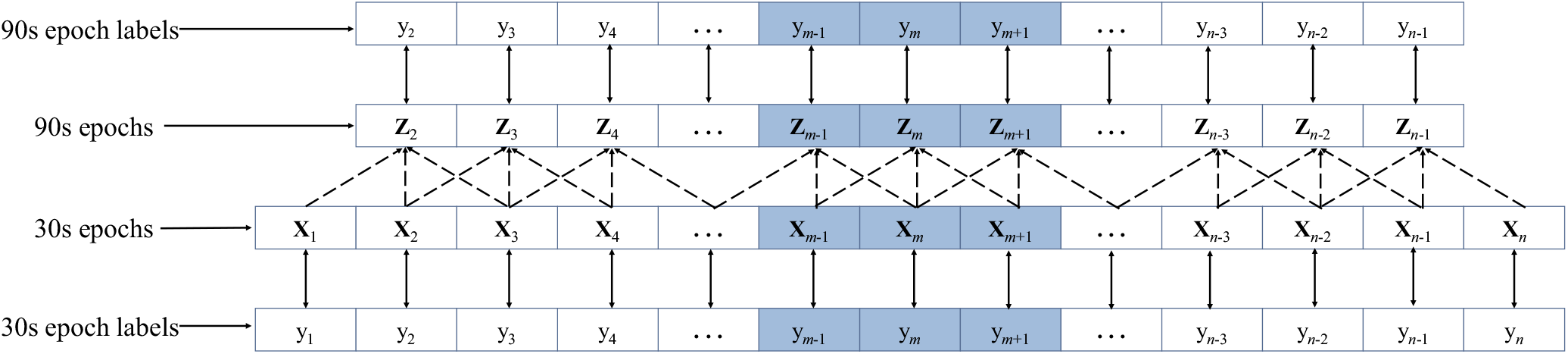
Illustration of 90s epochs and labels used in this paper, n donates the number of 30s epochs for a subject, **Z**_***m***_ is comprised of **X**_***m−*1**_, **X**_***m***_ and **X**_***m*+1**_, **2 *≤ m ≤ n −* 1**.

**TABLE I.** Number of 90s Epochs for Each Sleep Stage from Experimental Datasets

| Dataset | W | N1 | N2 | N3 | REM | Total |
| --- | --- | --- | --- | --- | --- | --- |
| CCSHS | 211030 | 19221 | 249681 | 17242 | 110188 | 690372 |
| Sleep-EDF | 69518 | 21522 | 69132 | 13039 | 25835 | 199046 |
| Sleep-EDF-v1 | 10197 | 2804 | 17799 | 5703 | 7717 | 44220 |

## III. Proposed SCNet

Fig. 2 shows the overall architecture of the SCNet. The convolution block performs three operations sequentially: one-dimensional convolutional layer (Conv1D), batch normalization and M-Apooling1D. Similarly, each MC Block is followed by batch normalization, M-Apooling1D and Dropout layer in sequence. In our model, we employ the concatenation of max-pooling and average-pooling to take place of the maxpooling for capturing more representable features. Similar to the inception module [39], the MC block contains different sizes of convolutional filters to capture the corresponding information. Besides, we use the GAP layer to replace the traditional fully connected layer, and it is proved to be more robust spatial translations of the input without parameter optimization [40].

**Fig. 2.**
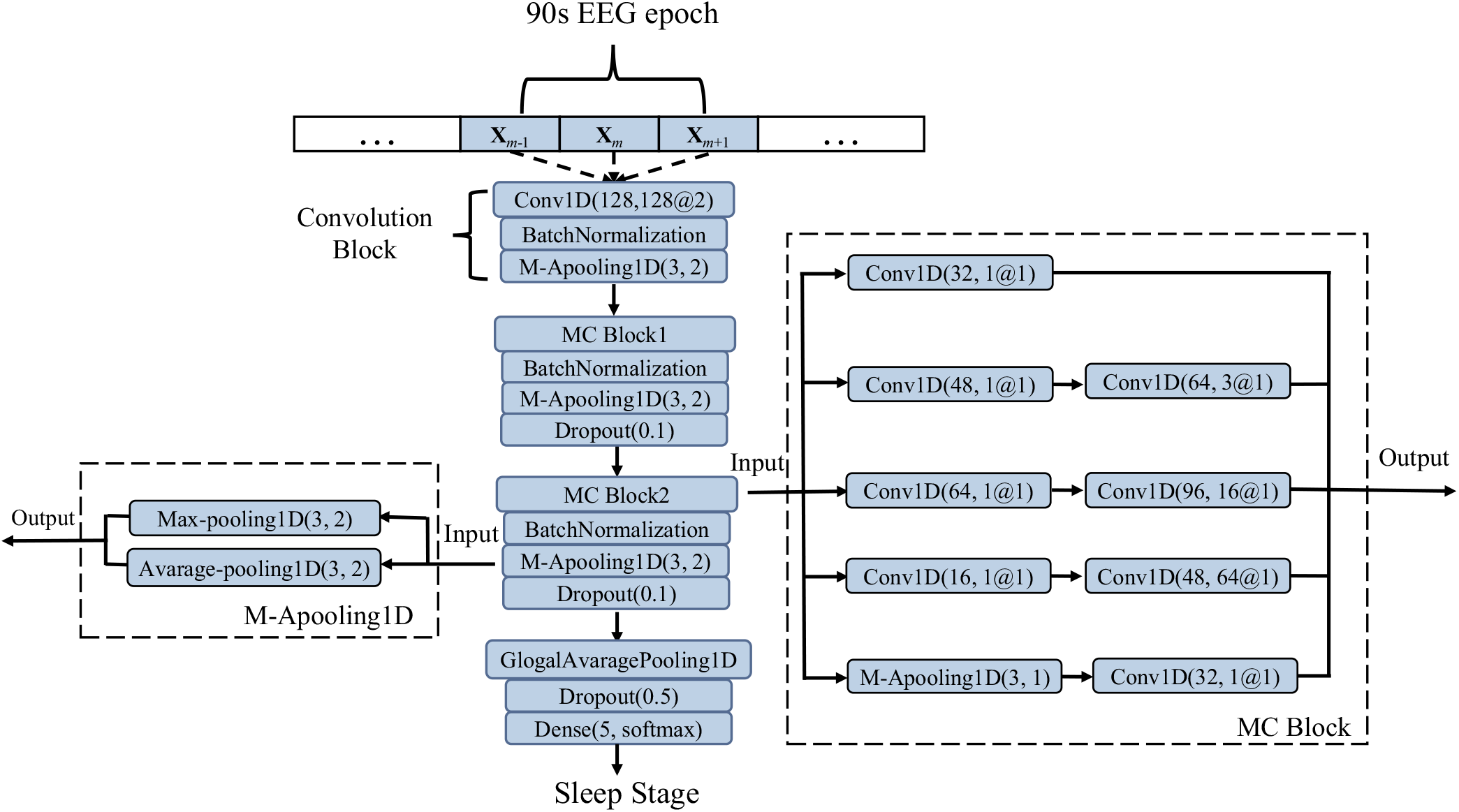
An overall architecture of the proposed SCNet.

### A. Model Specification

In Table II, we relate detailed parameters of the proposed model. The size of the model’s input is (90 *× f*_*s*_, 1), where *f*_*s*_ is the sampling rate. To be specific, the *f*_*s*_ of EEG C4 and Fpz-Cz is 128 Hz and 100 Hz, respectively. Here, the SCNet does not restrict the length of input which can be applied to different datasets.

**TABLE II.** Parameters of the Proposed Model

| Layer | Layer Type | Filters | Size | Stride | Activation | Output dimension |
| --- | --- | --- | --- | --- | --- | --- |
| 1 | Input | - | - | - | - | $(90 \times f_s, 1)$ |
| 2 | Conv1D | 128 | 128 | 2 | Relu | $(45 \times f_s, 128)$ |
| 3 | M-Apooling1D | - | 3 | 2 | - | $(\lceil 45 \times f_s/2 \rceil, 256)$ |
| 4 | MC Block1 | - | - | 1 | Relu | $(\lceil 45 \times f_s/2 \rceil, 272)$ |
| 5 | M-Apooling1D | - | 3 | 2 | - | $(\lceil 45 \times f_s/4 \rceil, 544)$ |
| 6 | Dropout(0.1) | - | - | - | - | $(\lceil 45 \times f_s/4 \rceil, 544)$ |
| 7 | MC Block2 | - | - | 1 | Relu | $(\lceil 45 \times f_s/4 \rceil, 272)$ |
| 8 | M-Apooling1D | - | 3 | 2 | - | $(\lceil 45 \times f_s/8 \rceil, 544)$ |
| 9 | Dropout(0.1) | - | - | - | - | $(\lceil 45 \times f_s/8 \rceil, 544)$ |
| 10 | GAP | - | - | - | - | 544 |
| 11 | Dropout(0.5) | - | - | - | - | 544 |
| 12 | Dense | - | - | - | Softmax | 5 |

The first convolutional layer with 128 filters of size 128 and a stride of 2 is applied to obtain the feature map from raw single-channel EEG. The activation function of this layer is rectified linear unit (ReLU) which is defined as the positive part of its argument:

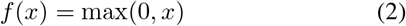

where *x* is the input of a neuron. To normalize the prior layer output, we apply the batch normalization technique. Besides, the M-Apooling layer can get the combination of maximum and average values from each of a cluster of neurons at the previous layer.

We implement two MC blocks in our model, and the filter sizes are selected among 1, 3, 16 and 64 to obtain multiscale representative features. More specifically, the small filter is prone to learn temporal information, while the large filter is better to capture frequency information. Considering the long length of input (128 × 90, 100 × 90), we optimize the filter sizes from the small sizes (3, 5 and 7), medium sizes (16 and 32) and big sizes (64, 128 and 256). The filter size of 1 is to improve the nonlinearity of the network and reduce the dimension of previous layer output. The filter sizes are chosen with 1, 3, 16 and 64 based on the optimized results. Furthermore, after concatenating the output of all convolutional layers, the dimension of the MC block1 output is (45 *× f*_*s*_*/*2, 272). The following M-Apooling layer can get (45 *× f*_*s*_*/*4, 544) dimension feature map. Each MC block is followed by a batch normalization layer, a M-Apooling layer with size of 3 and a dropout layer with the probability of 0.1. To find appropriate strides, we test 4 strides: 1, 2, 3 and 5. The stride of two MC blocks is set to 1, while the stride of the M-Apooling layer and the first convolutional layer is 2. The GAP layer is applied to flat the previous output before the final decision layer. Through a drop layer with drop rate of 0.5, the dense layer using softmax as the activation function makes the final decision. Softmax function can calculate the probabilities of five stages, the stage with maximum probability is as the consequence of the predicted sleep stage.

### B. Regularization

We adopt two regularization approaches to help prevent the overfitting problem. The first technique is L2 regularization that adds squared magnitude of coefficient as penalty term to the loss function. It is important to choose a proper regularization rate (lambda), if lambda is very large, it would add too much weight causing an underfitting issue. By contrast, a very small lambda would make the model more complex, then the model would learn too much about the particularities of the training data, L2 regularization therefore has little effect on avoiding overfitting. Hence, we test four lambda values: 10^*−*1^, 10^*−*2^, 10^*−*3^ and 10^*−*4^, the results show that 10^*−*3^ achieves the best performance. The L2 regularization is applied to all convolutional layers, including the MC block.

Another regularization method is dropout, which randomly drops units from the model during training with a specific probability from 0 to 1. It is noteworthy that the dropout layers is not used for testing. Dropout layers with probability of 0.1 and 0.5 are employed for the MC block and GAP layer, respectively.

### C. Training Setup

We select Adam as the network optimizer whose parameters ((learning rate) *lr, beta*1 and *beta*2) are set to 10^*−*3^, 0.9 and 0.999 respectively. Moreover, ReduceLROnPlateau of Callback in Keras is implemented to reduce the *lr*. Specifically, when the model monitors the validation accuracy showing no improvement within 3 epochs, the *lr* would drop to half of it. The minimum *lr* is set to 10^*−*7^. To find out appropriate batch size of mini-batch, size of 32, 64, 128, and 256 are evaluated, we select 64 as the size of mini-batch finally. The categorical cross entropy is chosen as the loss function of the model which is always used for classifying multi-class tasks. The model converges to the optimal solution within 40 iterations, hence the number of iteration is set to 40.

There are two types of methods to split the training and test sets [30], [41]. One is the subject-wise scheme which splits the training and test datasets based on the subjects. Another one is the epoch-wise method in which the split is conducted by epochs rather than subjects. In the epoch-wise scheme, We use 20% of whole data set as the test set and the remaining 90s epochs as the training set. As for the subject-wise approach, 80% subjects are selected as the training set, the other 20% subjects are used as the test set. Furthermore, we use the 5-fold cross-validation (80% training set for training, 20% training set for validation) scheme to train and evaluate our model for both datasets. In addition, only 90s epochs from the CCSHS dataset are used to determine the hyper-parameters of the proposed model. Once achieving optimal hyper-parameters, they would be used in all experiments. To be specific, when the model is applied to another dataset, there would be no need to modify the architecture and hyper-parameters of the model except for the input length which should adapt to the *f*_*s*_ of EEG from different datasets.

Graphic card Nvidia Tesla P100 with 16 Gbytes memory is used for model training. The implementation is written in Keras [42] with the Tensorflow backend [43].

## IV. Experimental Results

### A. Performance Metrics

We evaluate the model performance (epoch-wise) using accuracy (*ACC*), precision (*PR*), recall (*RE*), F1 score (*F* 1), and Cohen’s kappa coefficient (*K*). *ACC* is the proportion of correct predictions made by the model to the total predications. *PR* calculates the ratio of correctly predicted positives to all positives. *RE* means the fraction between true positives and all predications in the actual class. *F* 1 represents the weighted average of *PR* and *RE. K* measures the agreement between true labels and predicted labels. A large value of *K* can indicate good performance of the model. They are calculated as follows:

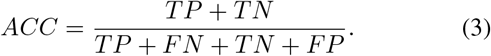

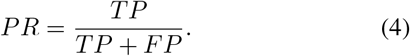

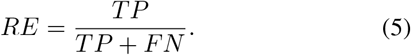

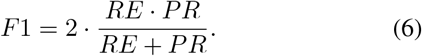

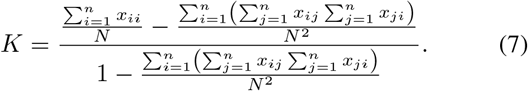

where *TP, TN, FN* and *FP* donate the true positives, true negatives, false negatives and false positives, respectively. *N* is the number of 90s epochs of the test set, *n* represents the number of classes. In this work, *n* equals 5, *x*_*ii*_ (1*≤ i ≤* 5) represents the diagonal value of the confusion matrix.

To show the performance of each fold cross-validation from the CCSHS and Sleep-EDF datasets, we present the normalized confusion matrices (CM) in Fig. 3. Firstly, we use single-channel EEG C4/A1 (90s epochs) from the CCSHS dataset to tune the hyper-parameters. Once getting the best performance, the hyper-parameters and model architecture are fixed for all experiments. Table III provides the mean CM of 5-fold cross-validation from the CCSHS dataset, we can see that the overall accuracy and *K* are respectively 90.2% and 86.5%. The proposed model shows the best ability to detect the W stage with the *PR* of 94.7%. By contrast, the performance of stage N1 classification is the worst which is consistent with the results of existing works. To be specific, there are 33.0% of N1 90s epochs being recognized correctly. In addition, 27.4% of N1 samples are misclassified as W, 19.5% as N2 and 20.1% as REM. Stages N2, N3 and REM have similar classification results in terms of the *PR* corresponding to 88.8%, 91.0% and 87.9% respectively.

**Fig. 3.**
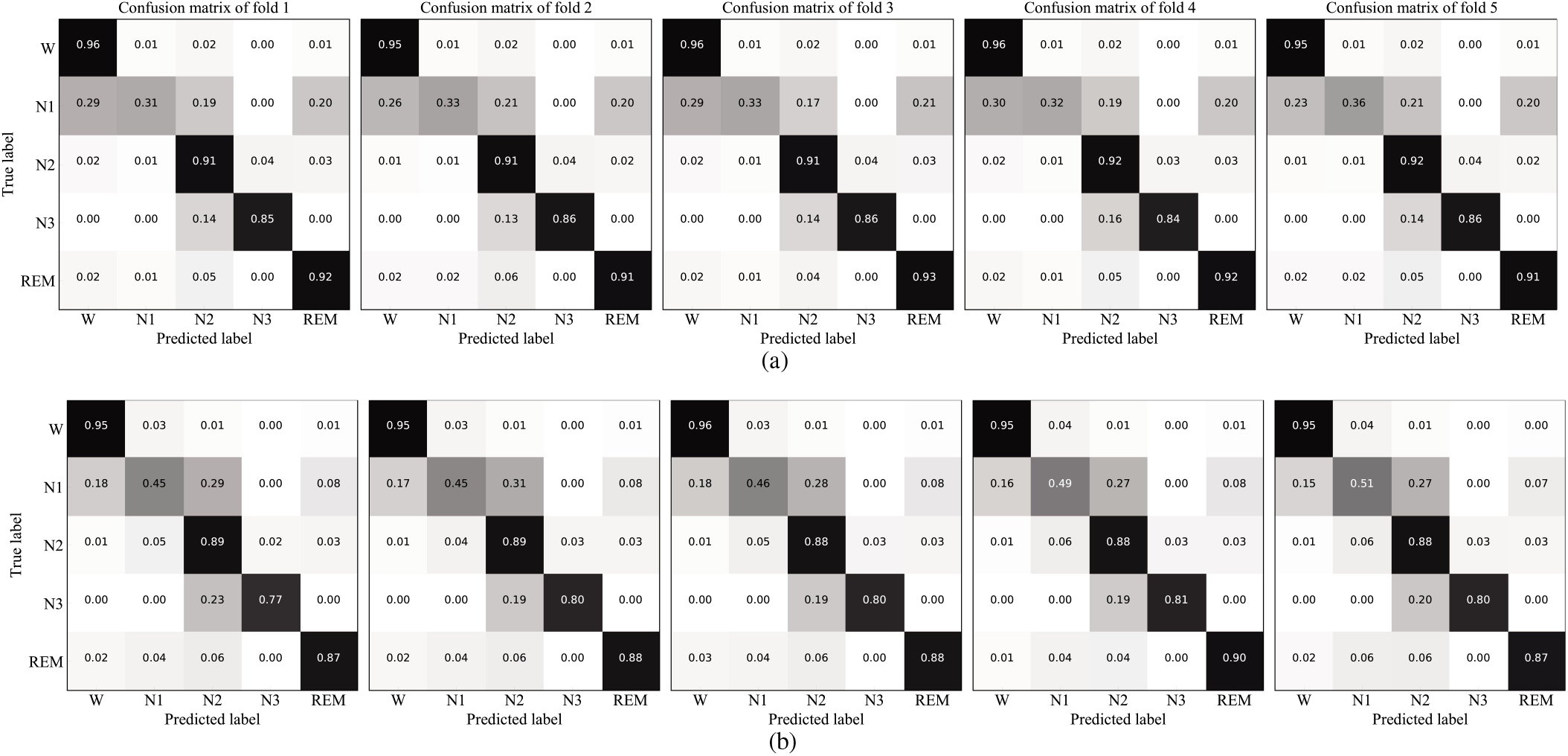
The normalized confusion matrices of each fold cross-validation. (a) CCSHS dataset and (b) Sleep-EDF dataset.

**TABLE III.** Mean Confusion Matrix of 5-Fold Cross-validation on Raw Single-channel EEG C4/A1 from the CCSHS Dataset

|  | Predicted |  |  |  |  | Per-class Metrics |  |  | Overall Metrics |  |
| --- | --- | --- | --- | --- | --- | --- | --- | --- | --- | --- |
| | W | N1 | N2 | N3 | REM | $PR(\%)$ | $RE(\%)$ | $F1(\%)$ | $ACC(\%)$ | $K(\%)$ |
| W | 40450 | 440 | 934 | 124 | 392 | **94.7** | **95.5** | **95.1** | 90.2 | 86.5 |
| N1 | 1039 | 1253 | 740 | 1 | 765 | 52.8 | 33.0 | 40.6 |  |  |
| N2 | 766 | 382 | 45679 | 1739 | 1359 | 88.8 | 91.5 | 90.1 |  |  |
| N3 | 60 | 0 | 3126 | 18791 | 8 | 91.0 | 85.5 | 88.1 |  |  |
| REM | 384 | 299 | 971 | 6 | 18358 | 87.9 | 91.7 | 89.8 |  |  |

To demonstrate the generalization capability of the proposed architecture, we also conduct the 5-fold cross-validation using the same model determined by the CCSHS dataset (i.e., without any hyper-parameters modification except for the input length) on the Sleep-EDF dataset. As can be seen from Table I, the distribution of the numbers of five stages is a bit different. Stage W has the biggest proportion and the number of N3 is the smallest in Sleep-EDF dataset, whereas the largest percentage is stage N2 in the CCSHS dataset. Besides, the EEG channel used in two datasets is also distinct, C4/A1 for the CCSHS dataset and Fpz-Cz for the Sleep-EDF dataset. It is worthy to note that despite the EEG channel and the size of the input length (90 *× f*_*s*_, 1) are quite different, the proposed model can obtain promising performance on two different datasets by comparing Table III and Table IV.

**TABLE IV.** Mean Confusion Matrix of 5-Fold Cross-validation on Raw Single-channel EEG Fpz-Cz from the Sleep-EDF Dataset

|  | Predicted |  |  |  |  | Per-class Metrics |  |  | Overall Metrics |  |
| --- | --- | --- | --- | --- | --- | --- | --- | --- | --- | --- |
| | W | N1 | N2 | N3 | REM | $PR(\%)$ | $RE(\%)$ | $F1(\%)$ | $ACC(\%)$ | $K(\%)$ |
| W | 14650 | 498 | 142 | 8 | 82 | **94.1** | **95.3** | **94.7** | 86.1 | 80.5 |
| N1 | 712 | 2020 | 1214 | 11 | 330 | 58.4 | 47.1 | 52.1 |  |  |
| N2 | 109 | 705 | 12255 | 385 | 383 | 84.9 | 88.6 | 86.7 |  |  |
| N3 | 4 | 5 | 532 | 2127 | 4 | 84.0 | 79.6 | 81.7 |  |  |
| REM | 99 | 233 | 299 | 1 | 4573 | 85.1 | 87.9 | 86.5 |  |  |

We further reveal the hypnogram comparison labeled by experts and the model’s prediction for one subject of CCSHS and Sleep-EDF datasets in Fig. 4.

**Fig. 4.**
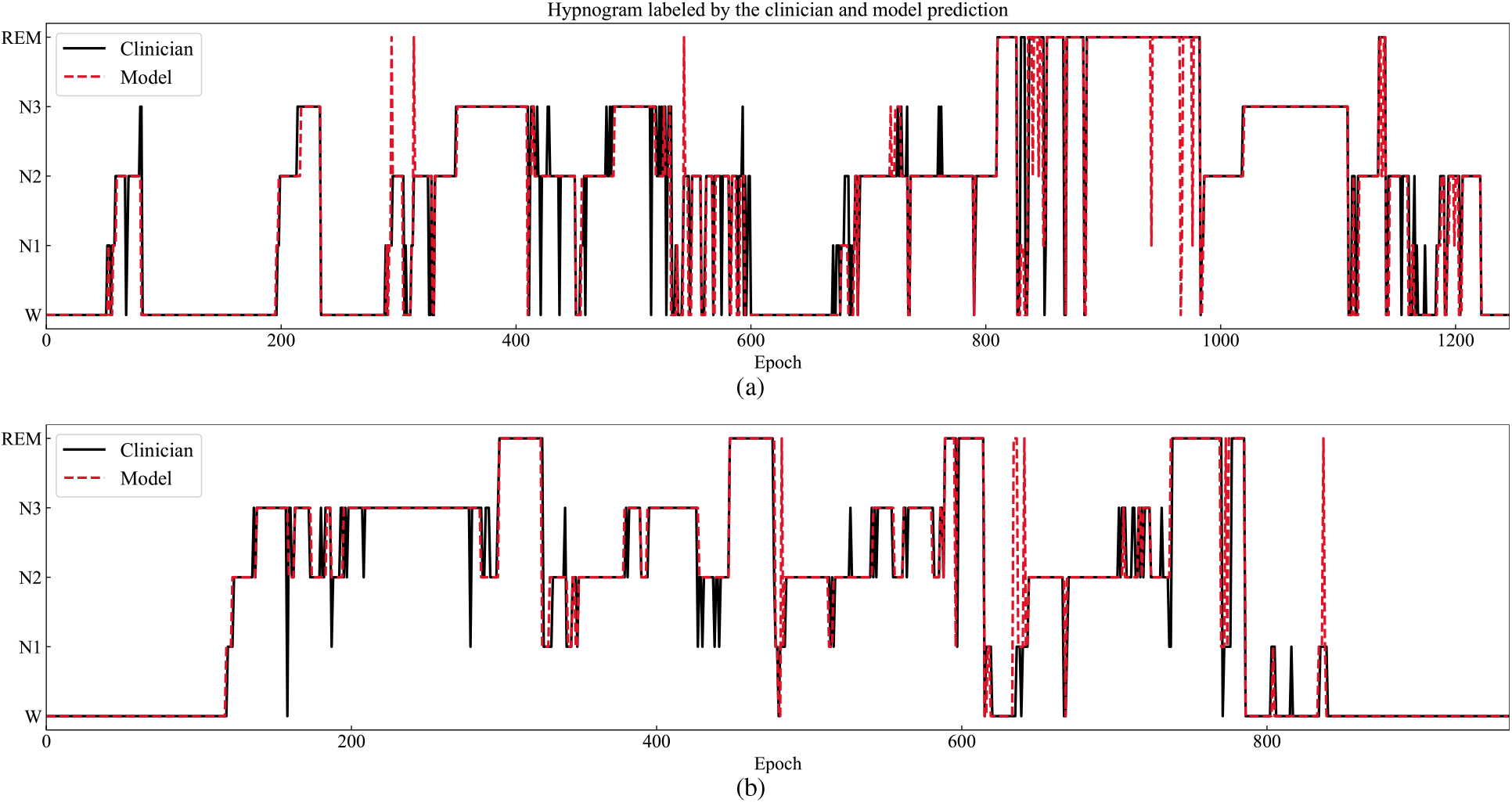
The comparison between hypnogram labeled by the clinician and the model’s prediction. The solid black line is the ground truth, the dotted red line donates the hypnogram labeled by the prediction of the proposed model. (a) CCSHS dataset and (b) Sleep-EDF dataset.

### B. Performance Comparison

We make a comparison between the proposed model (epoch-wise and subject-wise) with some existing works using the same datasets in terms of the *ACC* and *K* in Table V.

**TABLE V.**
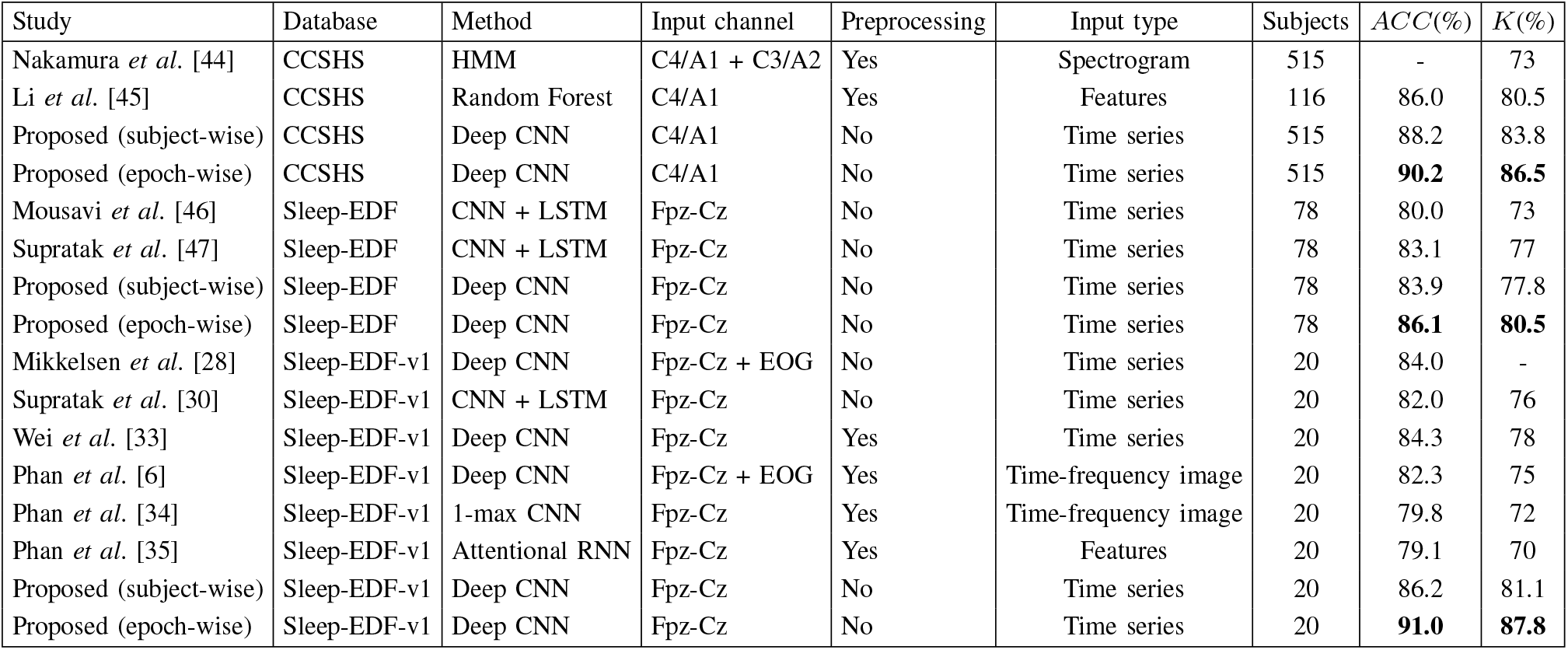
Performance Comparison between The Proposed Method and Previous Methods on the CCSHS, Sleep-EDF and Sleep-EDF-v1 Datasets

Table V reveals that the proposed framework can achieve higher *ACC* and *K* using raw single-channel C4/A1 EEG compared to approaches using multi-channel PSG data [44] or the single-channel EEG [45] on the CCSHS dataset. For the Sleep-EDF and Sleep-EDF-v1 databases, the proposed model also achieves comparable performance compared to state-of-the-art methods. Some studies [34], [35] extract features manually or multi-channel signals are used as input [6], [28] or some methods adopt single-channel EEG [33], [34], [46], [47]. Considering results of the comparison, the proposed framework can achieve promising performance on CCSHS, Sleep-EDF and Sleep-EDF-v1 datasets.

## V. Discussion and Conclusion

In this paper, we propose an end-to-end framework with CNNs, namely SCNet, which combines the feature learning ability and classification capacity. The proposed model is applied to classify sleep stages automatically from raw single-channel EEG without using any hand-engineered features and any other preprocessing (e.g., signal filtering and resample implementation). There are two main advantages that we train and evaluate the model with raw single-channel EEG. Comparing with those methods with hand-crafted features [4], [12], [48], where extracting hand-engineered features is conducted with priori knowledge and not in a data-driven way, and it is time-consuming for the researchers. Moreover, the selection of types and number of features would result in different model performance, there is no gold standard about the extraction of hand-crafted features. The second advantage is that it is much easier and more comfortable to record single-channel EEG data compared to the multi-channel scheme [6], [28] either at the hospital or home. Moreover, multi-channel PSG data used as input can increase the computational cost. Considering practical applications, the use of raw single-channel EEG can simplify the measurement scheme and reduce the related cost.

Comparing with the conventional deep neural network based on CNNs, where the convolutional layers with the fixed filter size are assembled in sequence. In such a case, it is not capable of capturing features representation from different scales. To address this issue, our model employs two MC blocks, which are the concatenation of several convolutional layers with four distinct filer sizes, to extract different scale features. Instead of using the traditional max-pooling layer, we adopt the M-Apooing layer to add average feature representation with maximum features simultaneously, which further improve the proposed model’s ability of feature learning. In addition, the SCNet model is quite simple and compact with a total 5 × 10^5^ parameters compared to the methods in [46] which has 2.1 × 10^7^ parameters and [30] in which the number of parameters of the representation learning and sequence residual learning parts has up to 6 × 10^5^ and 2 × 10^7^ respectively. Moreover, the proposed SCNet model can achieve the comparable performance with less computing resources occupied. Concerning online and realtime applications (e.g., sleep monitoring), our model with raw single-channel EEG is more reasonable to reduce the time latency and obtain reliable results.

To demonstrate the generalization of the proposed architecture, different single-channel EEGs from two datasets are adopted. The length of input is not restricted to a fixed number, our model can be adapted to different length of input relating to the *f*_*s*_ of EEG efficiently. Experimental results show that the proposed model can obtain promising performance on two datasets (CCSHS: *ACC*-90.2%, *K*-86.5%; Sleep-EDF: *ACC*-86.1%, *K*-80.5%), which indicate the desirable generalization of the SCNet model.

It is challenging to train on dataset A and test on B, not only for the proposed SCNet but also for typical CNNs. CNNs are running in a data-driven way which means the model must learn some crucial features from the training samples. Otherwise, it cannot perform well on an unfamiliar dataset. This is also the biggest difference (generalization ability) between machine and human, human beings are good at deducing and inducing. To further show the generalization ability of the proposed model, we perform two additional experiments. Firstly, we train our model with the CCSHS database, the obtained model then is tested on the Sleep-EDF dataset without any training, the accuracy is 65.9%. In reverse, The proposed model is trained on the Sleep-EDF dataset and tested with the CCSHS database, the accuracy achieved is 70.2%. In our future work, we will try to construct a more brain-inspired model with some cognitive neural dynamic from neuroscience [49] to increase the generalization ability for the sleep stage classification task. Also, it is valuable to adopt clinic datasets that have rarely been explored in previous studies.

## Acknowledgment

This study is to memorize Prof. Tapani Ristaniemi from University of Jyväskylä for his great help to the authors and Prof. Tapani Ristaniemi has supervised this study very much.

